# Biome-specific genome catalogues reveal functional potential of shallow sequencing

**DOI:** 10.1101/2025.06.16.659887

**Authors:** Alejandra Escobar-Zepeda, Matti O. Ruuskanen, Martin Beracochea, Jennifer Lu, Dattatray Mongad, Lorna Richardson, Robert D. Finn, Leo Lahti

## Abstract

The use of 16S rRNA metabarcoding for functional prediction is limited by several biases. Shallow shotgun sequencing is a cost-effective and taxonomically high-resolution alternative to 16S rRNA metabarcoding, but the low sequencing depth limits functional inference. Our BioSIFTR tool maps shallow shotgun sequencing reads against single-biome databases and extrapolates into their precalculated functional profiles. We used three datasets from red junglefowl, mice, and human gut, containing matched deep shotgun metagenomic and 16S rRNA metabarcoding data for taxonomic and functional benchmarking. An additional human gut deep shotgun sequencing data set was subsampled to 1 M reads and analysed with BioSIFTR to replicate previously obtained results. BioSIFTR taxonomic and functional profiles closely agree with the results of the full deep sequencing data in all biomes. We also replicated differences in the human gut microbiome between high and low trimethylamine N-oxide producing participants, using only < 2 % of the original deep sequencing data. The BioSIFTR tool is a powerful approach which approximates the functional information of a deep-sequenced metagenome while using only a fraction of the data. Shallow shotgun sequencing combined with BioSIFTR could be a stand-in replacement for 16S rRNA metabarcoding with an increased taxonomic and functional resolution, and lower bias.

**Graphical abstract:** 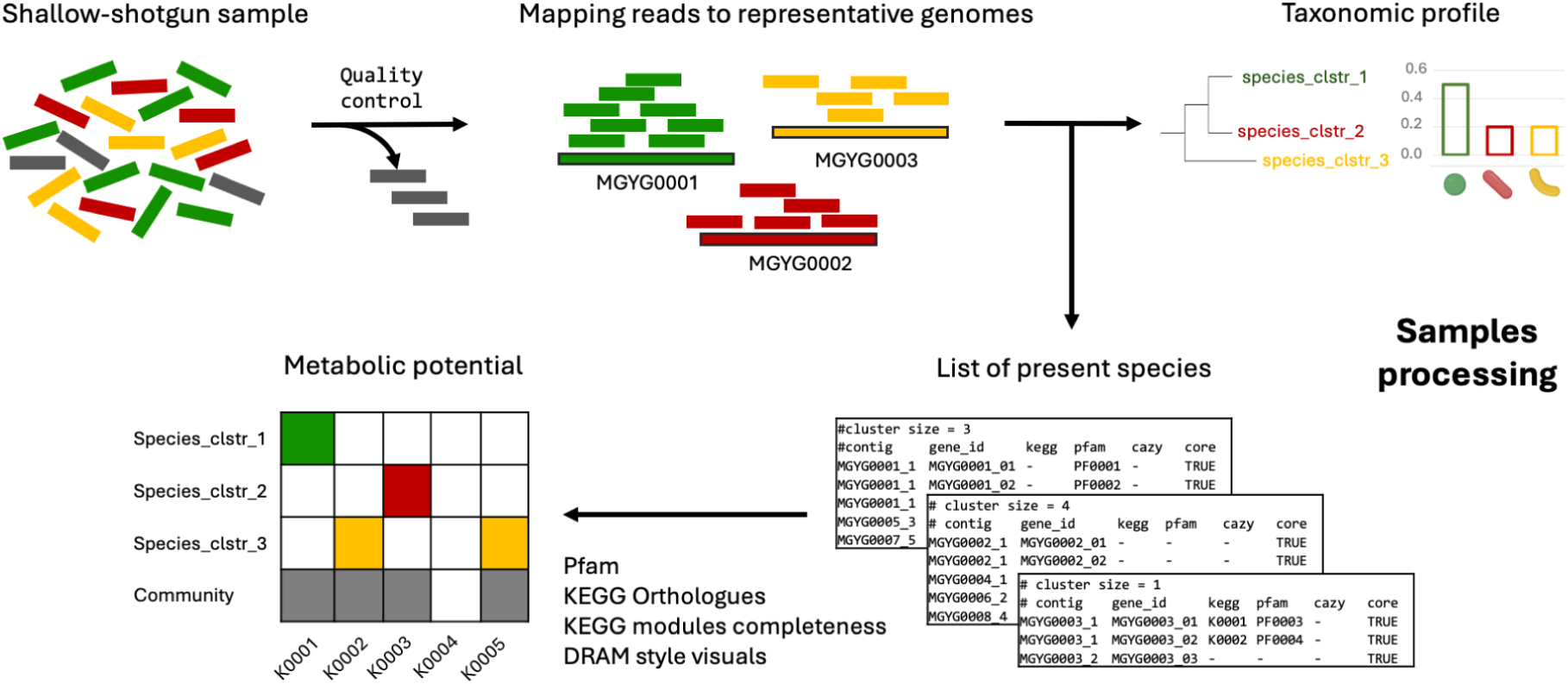

## 1. Introduction

The gold standard for determining the functional potential of a microbial community is the application of deep shotgun sequencing and subsequent transfer of functional annotation from databases. However, the high cost of sequencing, often to ∼50 million (M) reads per sample, can be prohibitive and further exacerbated when processing a large number of samples. A high number of samples is commonly required in a clinical setting, for spatial and longitudinal sampling, and in population studies. In other cases, low biomass or high host contamination can reduce the sequencing depth regardless of effort (1). Another issue can be the cost of computationally analysing these large datasets, with metagenomic assembly typically requiring access to high-memory (up to several TBs of RAM) computing resources (2). Thus, researchers face the tradeoff between a deep representation of the sample and the cost per run.

Currently, the low-cost option for studying microbial communities is via metabarcoding. This approach involves amplifying a specific marker gene, typically the small subunit ribosomal RNA (rRNA) gene (16S rRNA for Bacteria and Archaea, and 18S rRNA for Eukaryota) and sequencing up to ∼100,000 reads per sample (3). The resulting sequence reads are easily converted into a taxonomic profile, which can be used to extrapolate a functional profile (4, 5). However, such metabarcoding approaches, especially with short-read sequencing technologies, lack taxonomic resolution due to inherent limitations of the gene regions used as markers (6). Furthermore, the subsequent functional inference contains many sources of inaccuracies, given the aforementioned resolution and the lack of genomic references (7).

An alternative approach between deep-shotgun metagenomics and metabarcoding is a “shallow” shotgun metagenomic sequencing (up to ∼2 M reads per sample), which represents a viable option for low-cost functional inference. However, to our knowledge, few bioinformatic methods have been developed for functional predictions for shallow shotgun metagenomic sequencing data. Finding representative genomes and identifying low-abundant species have remained key challenges, leading to sparse and incomplete functional predictions. Currently, the most popular approach is mapping the reads directly to reference genomes or protein databases (8–11).

Using generalist databases for mapping metagenomic reads can easily lead to identifying organisms which are not a part of the studied microbiome, but rather e.g., temporary visitors or contaminants (12). Also, conducting read mapping to databases including draft genomes contaminated with human DNA can cause false positive assignments (13). Microbial species and genes tend to be biome-specific and form enrichment patterns through a biome–species–function relationship shaped by ecological and evolutionary principles (14). Strains of the same species found in different environments can exhibit vastly different functional profiles (15), while different species inhabiting the same biome can share similar functions due to their adaptation to the same environment (16). In most studies, mapping metagenomic reads to curated biome-specific databases could thus alleviate problems with false positive or biologically irrelevant taxonomic and functional assignments.

The primary goal of this work was to develop a robust bioinformatics solution for the interpretation of biome-specific microbial community functions from ’shallow shotgun’ metagenomes. To address this challenge, we implemented the *BioSIFTR* (Biome-specific Shallow shotgun Inference of Functional Traits through Read-mapping) pipeline to map raw reads from shallow shotgun sequencing to biome-specific microbial genome catalogues in MGnify (17), to identify the species present in the sample. Subsequently, a functional profile of the microbial community can be derived based on the pangenomes of the species identified. The tool is implemented in Nextflow and openly available (https://github.com/EBI-Metagenomics/BioSIFTR/).

In this work, we present the selection and parameter optimisation of mapping tools by evaluating their predictive capabilities on synthetic microbial communities, and using these results to develop the BioSIFTR pipeline. We then compared the pipeline’s performance on real data to the results from SHOGUN, a widely-used tool for shallow shotgun analysis (18). For completeness, we included functional inference comparisons using the 16S amplicon methods PICRUSt2 (4) and MicFunPred (5), alongside deep-shotgun assembly and functional annotation methodologies as the gold standard.

## 2. Methods

### 2.1. Mapping tools optimisation based on taxonomic prediction power

We benchmarked the performance of two different methods for evaluating the presence of species in metagenomic samples of human gut and chicken gut (see Supplementary material, Figure S1): the read-mapping tool bwa-mem2 v2.2 (19). and the k-mer content-based tool Sourmash v4.8.4 (20). To do so, we used a set of genomes, consisting of isolates, metagenomic assembled genomes (MAGs), and single-cell amplified genomes (SAGs), not included in the MGnify Genomes catalogues, to produce microbial synthetic communities with a variety of species richness. We selected representative genomes meeting the following quality control criteria: more than 50% complete, less than 5% contaminated, and having a minimum quality score of 50 (% completeness – 5 × % contamination) according to (17). For this benchmark, we used:

1. species in common with the MGnify Genomes to test taxonomic prediction power; and
2. the whole set of genomes to test functional prediction power to evaluate the tool’s capacity to detect functions even when the species is not present in the database.

The final human-gut dataset consisted of 1,623 genomes, of which 1,363 are NCBI isolates, 242 are MAGs/SAGs from PRJNA1030952, 11 are SAGs from PRJNA803937, and 7 are SAGs from PRJNA692334 (see accessions in the Supplementary material, Table S1). For the chicken-gut, we used the 2,625 representative MAGs genomes of the MetaChick project (21).

Subsequently, we generated synthetic Illumina paired-end raw reads using the InSilicoSeq tool v1.5.3 from the benchmark dataset. The microbial communities were designed to have species richness of 50 and 500 for the chicken gut, and 100 and 1,000 for the human gut. For each species richness level, we prepared 20 replicates with 1 M reads per replicate, resulting in a total of 160 sequence sets. Synthetic raw reads are available at the European Nucleotide Archive under the project accession PRJEB89162.

#### Statistical analyses for tool optimisation

To test the taxonomic prediction power, the raw reads were mapped to the corresponding MGnify Genomes catalogue using bwa-mem2 and filtered to keep reads with coverage > 90% and nucleotide identity > 60%. We then classified the mapped reads as all, best, and unique matches. To control for false positives, thresholds of mean coverage depth of 0, 1e-4, 1e-3, 1e-2, 1e-1, and 1 were tested. Sourmash performance was tested using sketches with k-mer sizes of 21, 31 (default), and 51.

The precision, recall and F1-score of the taxonomic predictions were calculated using the methods as described in (22), which takes into account the hierarchical nature of taxonomic annotations. In addition, False Positive (FP) and False Negative (FN) were counted and normalised by the number of True Positives (TP) per sample. Once the conditions for the taxonomic inference were optimised, we evaluated the prediction of functions in the different sets of synthetic communities.

### 2.2. Benchmark of functional inference

To produce the ground truth of the functional profile, each reference genome was annotated using EggNOG mapper v2.1.1 (23), and the resulting annotations were aggregated at the microbial community level. Annotations with e-values < 1e-10 were used to extract KEGG orthologues (KOs) (24) and Pfam annotations (25). The KEGG modules completeness was computed from the lists of unique KOs using the KEGG pathways completeness tool v1.0.1 (https://github.com/EBI-Metagenomics/kegg-pathways-completeness-tool). Pfam annotations assigned the “Type” Family were removed to avoid redundancy with lower hierarchy annotations. Performance metrics were calculated based on the presence/absence of the annotations using all the genes in the references (pangenome mode) or the core genes only (core mode). We computed statistical analysis and generated visuals using the ggbubr library v0.6.0 (26). Significance levels were added to the plots based on the Wilcoxon test (default significance threshold P < 0.05) without multiple testing.

### 2.3. Generation of functional annotation databases for the BioSIFTR pipeline

We generated EggNOG annotations of the MGnify Genomes catalogue for the pangenome genes for species with cluster size > 1. Subsequently, genes were labelled as core (true or false), and function tables were gathered for Pfam, KOs, and CAZy (27) annotations using an e-value cut-off of < 1e-10. KEGG modules’ completeness was calculated from the list of core or pangenome KOs with the KEGG pathways completeness tool. This set of functions represents the metabolic potential for the species. To calculate the metabolic potential of the community, the species representatives from the relevant catalogue are used as a reference database, and when a species is deemed to be present via read mapping, the functional profile of that species is transferred. The pipeline has the option to report core functions (conservative) or the functions observed in the entire pangenome.

### 2.4. Benchmarking BioSIFTR and other metagenomic approaches on real data

To evaluate the utility of our approach, we generated and compared the taxonomic and functional profiles on publicly available metagenomic samples of three gut biomes (see Supplementary material, Figure S2). The selected projects have deep shotgun metagenomic raw reads and 16S rRNA gene amplicon sequencing data from the same sample (i.e. sharing sample accessions SAMEA): human (PRJDB11444), mouse (PRJEB74255), and red junglefowl (PRJEB46806). We selected 30 pairs of samples from each project, ensuring a minimum of 20,000 reads for amplicon data and 20 M reads for shotgun metagenomic data. In addition, we randomly subsampled the deep shotgun data using the Seqtk v1.4 (https://github.com/lh3/seqtk) sample function to generate paired-end shallow shotgun raw reads with paired-end read counts of 0.1, 0.5, 1, 1.5, and 2 M.

The 16S rRNA gene amplicon data were processed in Qiime2 v2025.4 (28). After quality checking with the demux plugin v2025.4.0, reads were denoised to ASVs with the Qiime2 DADA2 plugin v2025.4.0 (29). PRJDB11444 reads were trimmed to 280 bp on the fw read and 220 bp on the rv read, PRJEB74255 reads were trimmed to 245 bp on the fw read and 220 bp on the rv read, and PRJEB46806 reads were trimmed to 230 bp on both fw and rv reads. Read pairing was done with ≥20 bp overlap in all datasets, and using the pooling method ‘pseudo’. Functionality was predicted first with MicFunPred v1.0.0, and then with PICRUSt2 v2.6.2, which uses the updated PICRUSt2-SC database (30). Taxonomy to the representative ASVs was assigned with the Qiime2 vsearch plugin v2025.4.0 (31) through the “non-v4-16” function in the Greengenes2 plugin v2024.1 (32), using the Greengenes2 database v2024.09. Data tables from Qiime2 outputs were exported with biom v2.1.14 (33).

Deep shotgun-sequenced raw reads were quality filtered using fastp v0.23.4 (34) and decontaminated by mapping to the reference genomes of human GRCh38 (GCF_000001405.40), phiX174 (NC_001422.1), and either chicken (GCF_016699485.2) or mouse (GCF_000001635.27) using bwa-mem2 v2.2.1 and samtools v1.10 (35). The high-quality reads were assembled using either SPAdes v3.15.5 (36) or Megahit v1.2.9 (37) if the reads required more than 200 GB of memory to assemble (see Supplementary material, Table S2). Contigs were used as input for functional annotation with EggNOG mapper. In addition, high-quality decontaminated raw reads were taxonomically annotated using the mOTUs pipeline v3.0.3 (38).

The shallow shotgun metagenomes were processed to run SHOGUN v1.0.8 using ‘--level species --function’ flags and the MGnify BioSIFTR pipeline v1.2.0 with the settings described in section 2.6 of this paper. Subsequently, the annotation results were integrated into matrices to enable comparative analysis. Taxonomic annotation tables were aggregated at different ranks and normalised by relative abundance. KEGG orthologues (KOs) annotation was processed as presence/absence tables.

#### Statistical analyses in comparisons against real data

A Jaccard dissimilarity matrix was constructed to determine the minimum number of reads that most closely resembled the gold-standard deep-shotgun KOs functional profiles. The BioSIFTR results from Sourmash and bwa-mem2, as well as SHOGUN subsampled results, were compared to the gold-standard deep-shotgun profiles. Statistical analysis was performed using Analysis of Similarities (ANOSIM) combined with a dispersion test (betadisper), utilising the vegan R package v2.6 (39). For pairwise dispersion analysis, we used a Tukey honest significant differences test (TukeyHSD) in R.

To compare with the amplicon-based method, we used the standard 1 M reads for SHOGUN and BioSIFTR results. Ordination plots were produced using the PCoA method in phyloseq R v1.42.0 (40) for a visual representation of group separation using Jaccard dissimilarity for KOs and Bray-Curtis dissimilarity for species.

### 2.5. Application of the BioSIFTR pipeline to a real case study

We tested our method on the detection of differentially abundant features in the phenotype differentiation of the human gut microbial conversion of L-carnitine into trimethylamine N-oxide (TMAO) (41). To do so, we retrieved the deep shotgun metagenomes corresponding to Healthy cohort 1 available in ENA under the project accession number PRJEB66719. We selected samples having a minimum of 20 M raw reads, and the reads were processed for quality control and host-decontaminated using the strategy described above. Then, we generated shallow shotgun data by randomly subsampling to 1 M raw reads and annotated using our BioSIFTR pipeline with the bwa-mem2 tool in ‘pan’ mode. We tested for significant differences between high and low TMAO producers, as well as before and after L-carnitine intervention, using both taxonomic (species rank) and functional (KOs and Pfam annotation) matrices.

#### Statistical analyses with case study data

Dissimilarity matrices were generated using Bray-Curtis for species relative abundance and for KOs and Pfam functions after normalisation by cumulative sum scaling. Then, global PERMANOVA (adonis2) analysis in combination with a dispersion test (betadisper) using the vegan R package v2.6. When significant differences between groups were observed, an analysis of similarities (ANOSIM) was computed to confirm the observation. The alpha or significance level was set at P < 0.01 across all methods. Differentially abundant functions were tested using MaAsLin 3 v0.99.16 (42) with the following parameters: max_significance = 0.1, normalization = TSS, correction = BH.

### 2.6. The MGnify BioSIFTR pipeline

#### Novel software

The BioSIFTR tool was developed using Nextflow and containerised tools, minimising barriers for installation and ensuring high reproducibility. Requirements include Nextflow version >=21.10 and container technology such as Docker (43) or Singularity (44). The code is publicly available on GitHub at https://github.com/EBI-Metagenomics/BioSIFTR/. The main modules of the pipeline are raw reads quality control, taxonomic profiling at species cluster level, transfer of the metabolic potential, and output writing (Figure 1).

**Figure 1.**
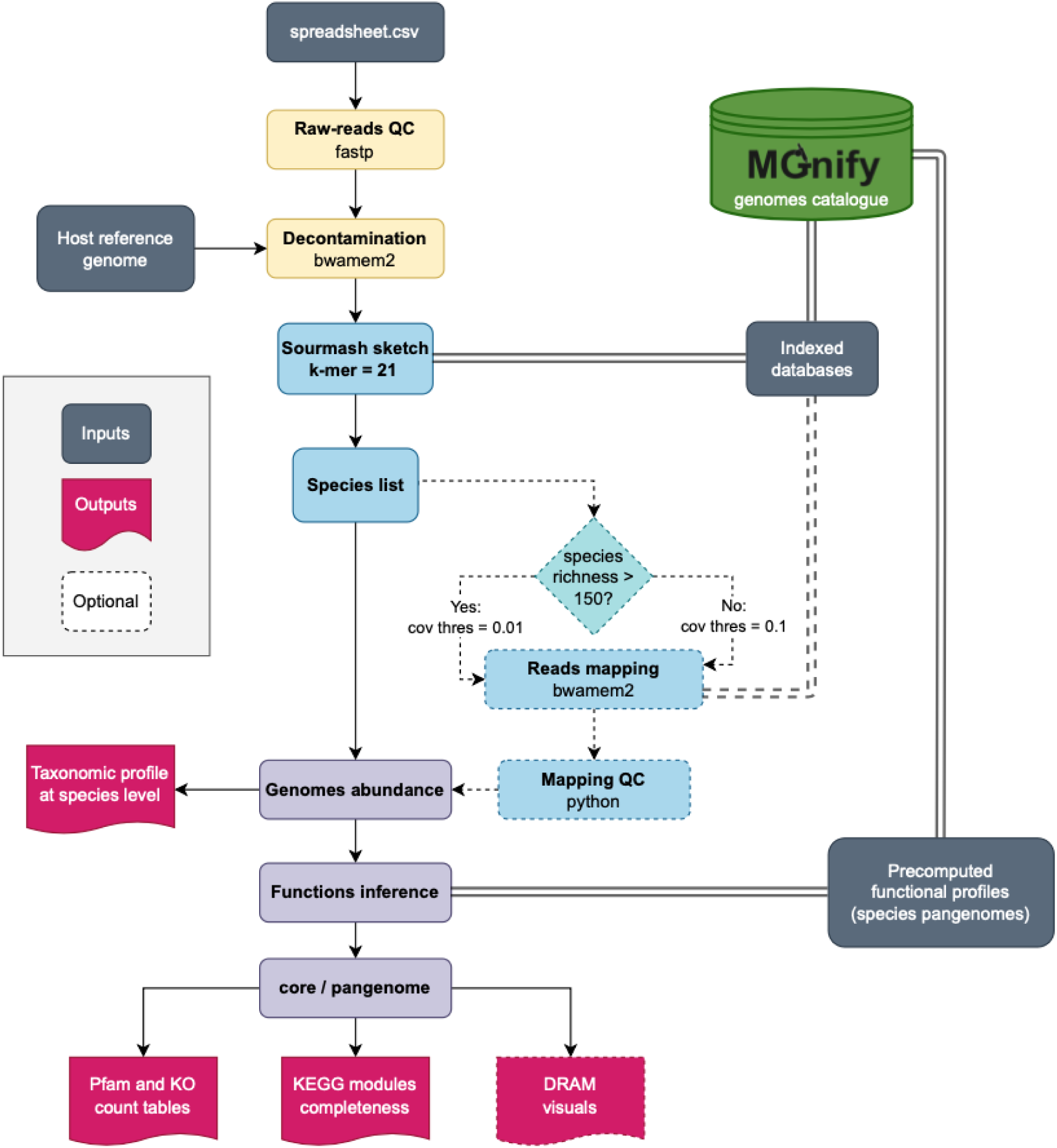
Overview of the MGnify BioSIFTR pipeline. Bwa-mem2 optimisation is dynamic.

The reference databases of representative species indexed for Sourmash and bwa-mem2, as well as the precomputed databases of functions per species cluster for all of the MGnify Genomes catalogues, are hosted by MGnify and are publicly available. Supported MGnify Genomes catalogues at the date of this publication are chicken-gut v1.0.1, cow-rumen v1.0.1, human-gut v2.0.2, human-oral v1.0.1, human-vaginal v1.0, honeybee-gut v1.0.1, marine v2.0, mouse-gut v1.0, non-model-fish-gut v2.0, pig-gut v1.0, sheep-rumen v1.0, and zebrafish-faecal v1.0. Databases are automatically downloaded the first time the BioSIFTR is run with a specific biome and can be updated with new versions. In addition, custom genome catalogues can be generated following the steps described in https://github.com/EBI-Metagenomics/biosiftr_extended_methods.

## 3. Results

### 3.1. Optimisation of tools for taxonomic prediction

We selected two popular methods to identify species presence and relative abundance in synthetic microbial communities: the read-mapping tool bwa-mem2 and the k-mer content-based Sourmash.

Performance metrics were calculated using the complete lineage information in the taxonomic predictions. The higher values of F1-score were observed using bwa-mem2 unique alignments. However, the optimal mean coverage threshold depends on the species richness of the dataset, with a coverage threshold of 0.1 for richness 50 and 100 and 0.01 for richness 500 and 1,000 (see Supplementary material, Table S3). In the same way, the optimal Sourmash k-mer settings differed according to species richness, with a k-mer of 51 for richness of 50 and 100; and a k-mer size of 21 for richness of 500 and 1,000.

We analysed the number of FP and FN in the two optimisation conditions for bwa-mem2 and Sourmash (Figure 2). In contrast, as the number of species increases in the microbial community, more species exceed the detection limit of the method, implying more false negatives.

**Figure 2.**
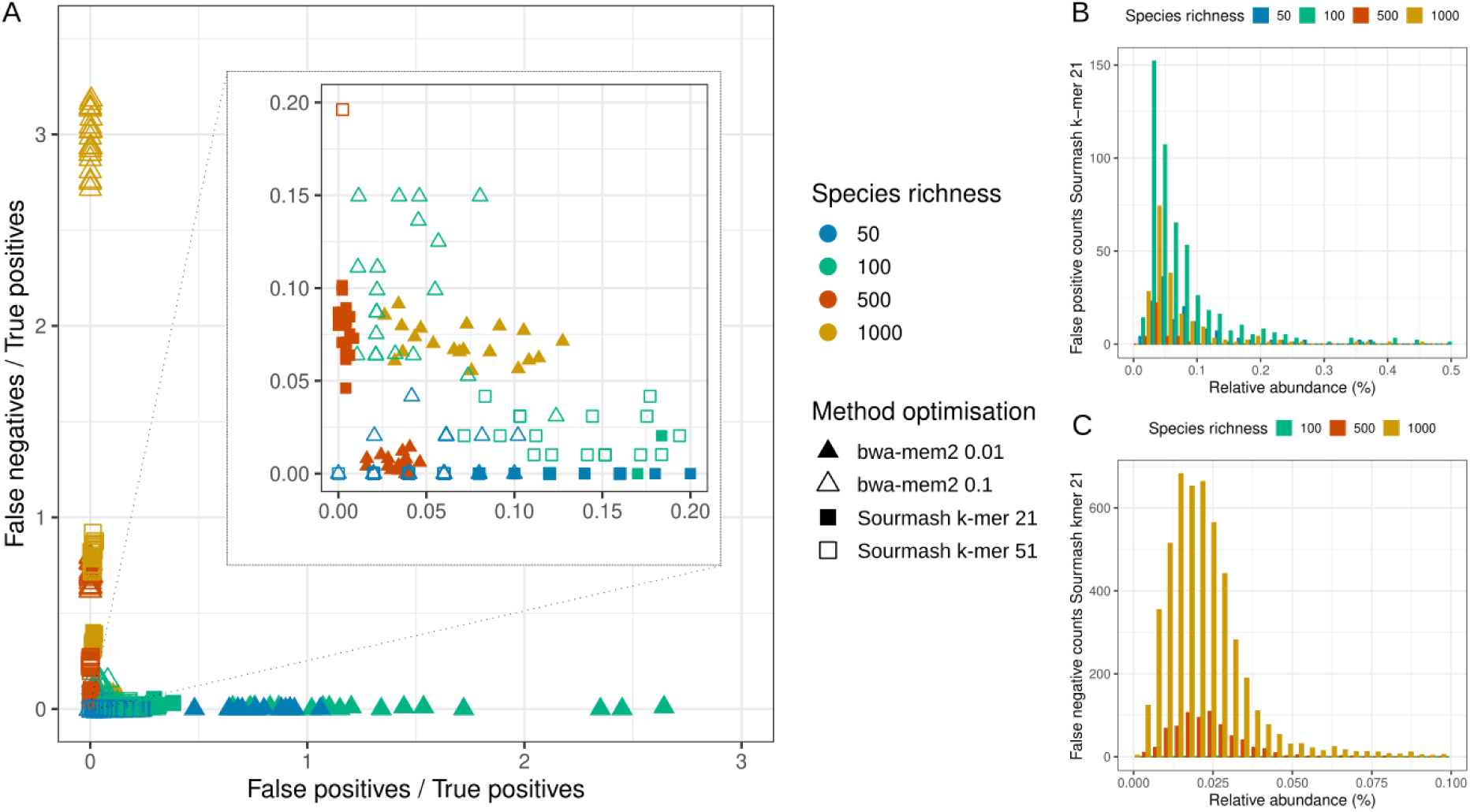
(**A**) Taxonomic prediction performance of BioSIFTR when using Sourmash and bwa-mem2 in terms of FP and FN counts at the species level. The optimal parameter of each tool depends on the microbial community species richness. A correct optimisation brings the data to the origin of the scatterplot. The distribution of relative abundance of (**B**) false positive and (**C**) false negative predictions using Sourmash kmer = 21.

We observed that the bwa-mem2 tool is more sensitive to suboptimal parameter choices than Sourmash. However, bwa-mem2 outperforms Sourmash with the proper optimisation. Using Sourmash with a k-mer size of 21 is optimal for species richness of 500 and 1,000, with a very low number of false positives. However, when the species richness is low, the method becomes prone to generating false positives. Also, it is important to note that the mean relative abundance of false negatives in the worst scenario is as high as 0.36 % for the 100-species richness human-gut dataset, and < 0.02% for the other datasets. This is the detection limit of the method. In addition, false positives occur with low frequency, adding species to the taxonomic profile with relative abundance < 0.071 % on average.

The optimisation of the tools for taxonomic prediction depends on the species richness of the microbial community. We selected Sourmash with k-mer size = 21 as the default mapping choice on the BioSIFTR pipeline because it is more robust than bwa-mem2 on samples with a low number of species. However, due to the good performance of bwa-mem2 with the optimal settings, we kept this tool as part of the pipeline under dynamic optimisation based on Sourmash species richness prediction. We set the coverage threshold to 0.01 for samples with species richness > 150 or 0.1 for species richness <= 150.

### 3.2. Functional inference on synthetic communities

Once the detection of the representative species cluster was optimised, we transferred the functional annotation of the species cluster as a contribution to the metabolic potential of the microbial community. The pipeline was designed to use the pangenome (pan) mode by default, but it can also be run with the annotation of core genes.

Based on the F1-score analysis, we observed distinct performance trends between pan and core modes across annotation types and between the two taxonomic prediction methods (Figure 3A and B). Globally, when a significant difference was observed, the pan mode consistently outperformed the core mode. The highest F1-scores were obtained for KEGG modules, highlighting the robustness of this type of annotation. Specifically, for Sourmash, the BioSIFTR in the pan mode showed superior performance over the core mode across all types of annotations, particularly for datasets with species richness greater than 100 (Figure 3A). In addition, KOs annotation consistently performed better in pan mode, regardless of species richness. Furthermore, when using bwa-mem2, a significant difference was seen between pan and core modes for KO and module annotations, but only for datasets with richness exceeding 50 species (Figure 3B). These findings highlight the advantages of pan mode, especially when working with larger, more diverse datasets.

**Figure 3.**
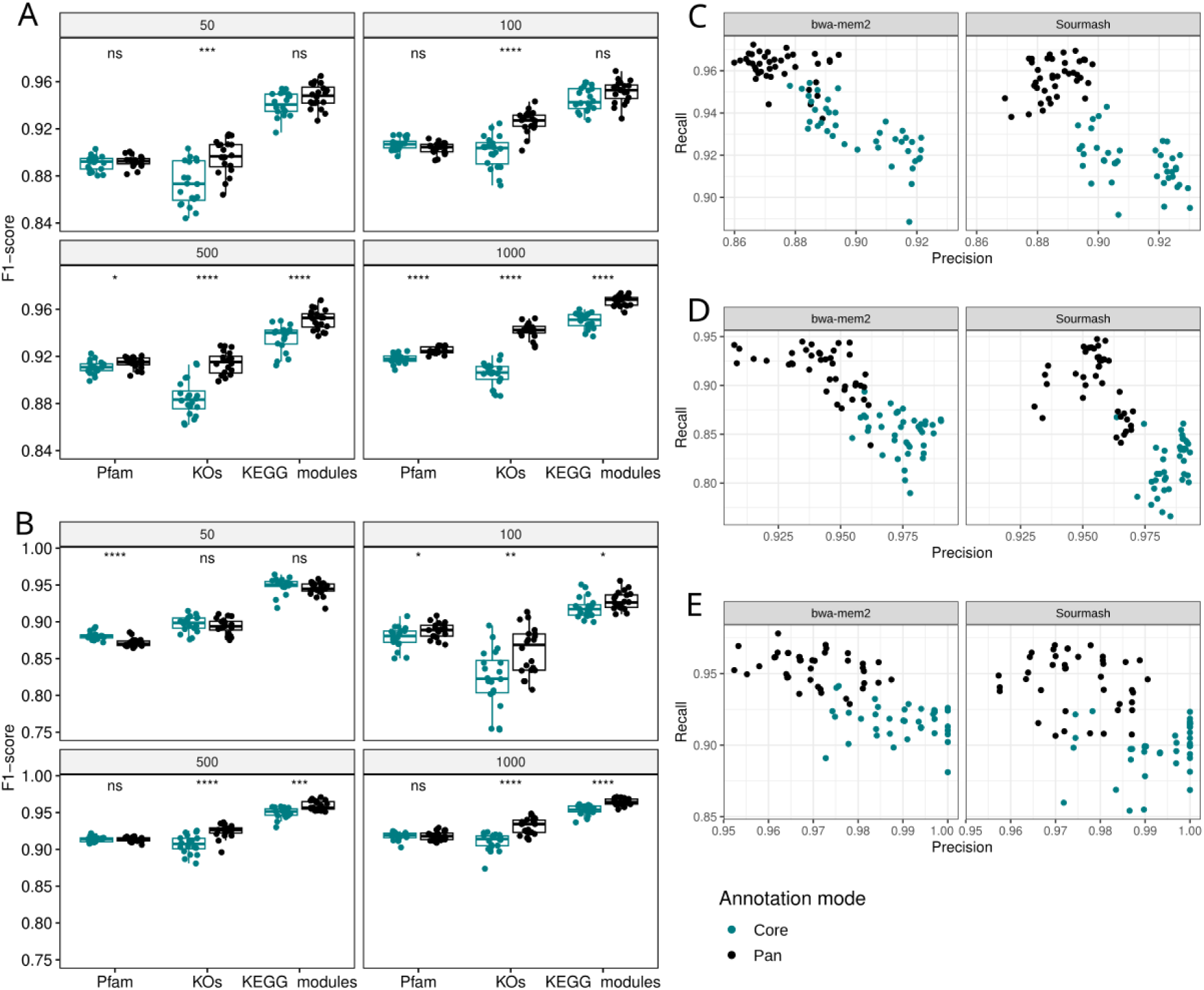
Functional inference performance based on taxonomy prediction as part of BioSIFTR tools optimisation. In panels (**A**) Sourmash k-mer size 21 and (**B**) bwa-mem2 dynamic optimisation, the F1-score performance metric of functional inference on synthetic communities is depicted. Significance corresponds to pairwise comparisons for the Wilcoxon test where * = 0.01 < p ≤ 0.05, ** = 0.001 < p ≤ 0.0, *** = 0.0001 < p ≤ 0.001, **** = p ≤ 0.0001. Right-hand panels show precision versus recall metrics on samples with higher species richness (500 and 1,000) for different types of functional annotation: (**C**) Pfam, (**D**) KEGG orthologues, and (**E**) KEGG modules.

The largest differences between the annotation modes (pan and core) were observed in samples with higher species richness (Figure 3A and B). In the core mode, BioSIFTR produced a lower number of expected functions because accessory genes are excluded, which negatively impacted the recall in the three types of functional annotation tested (Figure 3C-E). In contrast, while pan mode enhanced sensitivity, it also elevated the risk of overpredicting functions, leading to more false positives and a subsequent decline in precision. Nonetheless, the overall performance balance favoured BioSIFTR in the pan mode with both tools, bwa-mem2 and Sourmash.

### 3.4. Comparison of functional profiling approaches on real data

#### Selection of the shallowest sequencing from the subsampling experiment

We used the KO annotation results to find the most similar functional annotation versus its corresponding deep shotgun data as a gold standard reference.

The analysis of functional profiles generated from subsampled shotgun data of three biomes revealed significant differences in composition between the annotation tools and read depths (see Table 1). Regardless of the number of reads in the sample, SHOGUN consistently produced functional profiles that deviated from the gold standard. On the other hand, when the BioSIFTR pipeline was applied in combination with bwa-mem2, it generated functional profiles that closely resembled the gold standard in all three tested biomes, even at a subsampled level of 0.5 M reads in human and mouse gut. Note that dispersion is homogeneous between the BioSIFTR and deep shotgun profiles, meaning that differences are observed due to composition only.

**Table 1.** Comparative statistical analysis based on the predicted KOs functional profiles. Comparisons were made between the functional profiles of the gold standard deep-shotgun using Jaccard distance. R < 0.5, highlighted in bold, indicates profiles that are closely related to the gold standard. * lowest R-value per biome, indicating optimal performance.

| **Human-gut** | BioSIFTR Sourmash |  |  | BioSIFTR bwa-mem2 |  |  | SHOGUN |  |  |
| --- | --- | --- | --- | --- | --- | --- | --- | --- | --- |
| Subsampling (M) reads | P-adj (betadisp) | R-val (anosim) | P-val (anosim) | P-adj (betadisp) | R-val (anosim) | P-val (anosim) | P-adj (betadisp) | R-val (anosim) | P-val (anosim) |
| 0.1 | 0.999 | **0.296** | 0.001 | 0.914 | 0.909 | 0.001 | 0.260 | 0.996 | 0.001 |
| 0.5 | 0.603 | **0.320** | 0.001 | 0.998 | **0.340** | 0.001 | 1.000 | 0.953 | 0.001 |
| 1 | 0.117 | **0.397** | 0.001 | 1.000 | **0.292\*** | 0.001 | 0.998 | 0.956 | 0.001 |
| 1.5 | 0.009 | **0.463** | 0.001 | 1.000 | **0.320** | 0.001 | 0.999 | 0.944 | 0.001 |
| 2 | 0.004 | **0.496** | 0.001 | 1.000 | **0.388** | 0.001 | 0.928 | 0.943 | 0.001 |
| **Red junglefowl-gut** | BioSIFTR Sourmash |  |  | BioSIFTR bwa-mem2 |  |  | SHOGUN |  |  |
| Subsampling (M) reads | P-adj (betadisp) | R-val (anosim) | P-val (anosim) | P-adj (betadisp) | R-val (anosim) | P-val (anosim) | P-adj (betadisp) | R-val (anosim) | P-val (anosim) |
| 0.1 | 0.609 | 0.698 | 0.001 | 1.000 | 0.813 | 0.001 | 0.000 | 0.786 | 0.001 |
| 0.5 | 0.571 | **0.443** | 0.001 | 0.144 | 0.665 | 0.001 | 0.000 | 0.719 | 0.001 |
| 1 | 0.158 | **0.399** | 0.001 | 0.207 | **0.522** | 0.001 | 0.000 | 0.704 | 0.001 |
| 1.5 | 0.175 | **0.393** | 0.001 | 0.619 | **0.432** | 0.001 | 0.000 | 0.703 | 0.001 |
| 2 | 0.023 | **0.409** | 0.001 | 0.885 | **0.384\*** | 0.001 | 0.000 | 0.704 | 0.001 |
| **Mouse-gut** | BioSIFTR Sourmash |  |  | BioSIFTR bwa-mem2 |  |  | SHOGUN |  |  |
| Subsampling (M) reads | P-adj (betadisp) | R-val (anosim) | P-val (anosim) | P-adj (betadisp) | R-val (anosim) | P-val (anosim) | P-adj (betadisp) | R-val (anosim) | P-val (anosim) |
| 0.1 | 1.000 | **0.472** | 0.001 | 0.088 | 0.688 | 0.001 | 1.000 | 0.853 | 0.001 |
| 0.5 | 0.872 | 0.566 | 0.001 | 1.000 | **0.464** | 0.001 | 1.000 | 0.880 | 0.001 |
| 1 | 0.639 | 0.632 | 0.001 | 1.000 | **0.369\*** | 0.001 | 1.000 | 0.839 | 0.001 |
| 1.5 | 0.732 | 0.618 | 0.001 | 1.000 | **0.423** | 0.001 | 1.000 | 0.917 | 0.001 |
| 2 | 0.644 | 0.631 | 0.001 | 1.000 | 0.533 | 0.001 | 1.000 | 0.881 | 0.001 |

In the Junglefowl dataset, BioSIFTR’s performance improved progressively with increasing read depth (Table 1), suggesting that higher sequencing effort enhances functional resolution in less well-characterised biomes. In contrast, optimal performance for the mouse and human gut datasets was achieved at 1 million reads, likely reflecting the greater completeness and representation of these biomes in the reference database.

Comparisons in this section depend on the deep-shotgun assemblies’ quality (see Supplementary material, Table S2). Human-gut assemblies are of the highest quality (highest N50 values and assembled bases), making the comparison fairer. For this reason, and due to ∼ 0.5 - 2 M reads being the standard in shallow shotgun projects (10, 18, 45), we are using the results of subsampling to 1 M reads for the downstream analysis.

#### Shallow shotgun and 16S rRNA amplicon-based methods versus the gold standard Taxonomic annotation at species rank

The functional profiles generated by BioSIFTR are derived from the pangenomic functions of species clusters identified in the sample. When comparing taxonomic profiles obtained from deep shotgun sequencing (using the mOTUs pipeline, which annotates raw reads based on single-copy marker genes) with those generated by BioSIFTR, SHOGUN, and amplicon-based methods, significant compositional differences were observed across methods. However, data dispersion was homogeneous across the three tested biomes (see Supplementary material, Table S4), indicating that observed differences in taxonomic profiles are attributable to composition rather than variability. Among the tested approaches, BioSIFTR and deep shotgun sequencing produced the most similar taxonomic profiles across all three biomes (Figure 4). Note that amplicon-based profiles closely matched both deep shotgun and BioSIFTR results in the human-gut biome, but not in the other two, suggesting a database bias toward the human microbiome in amplicon-based annotations. Additionally, SHOGUN was the second-best-performing method in the mouse gut biome, following BioSIFTR, but produced more divergent profiles for the human gut and red junglefowl gut.

**Figure 4.**
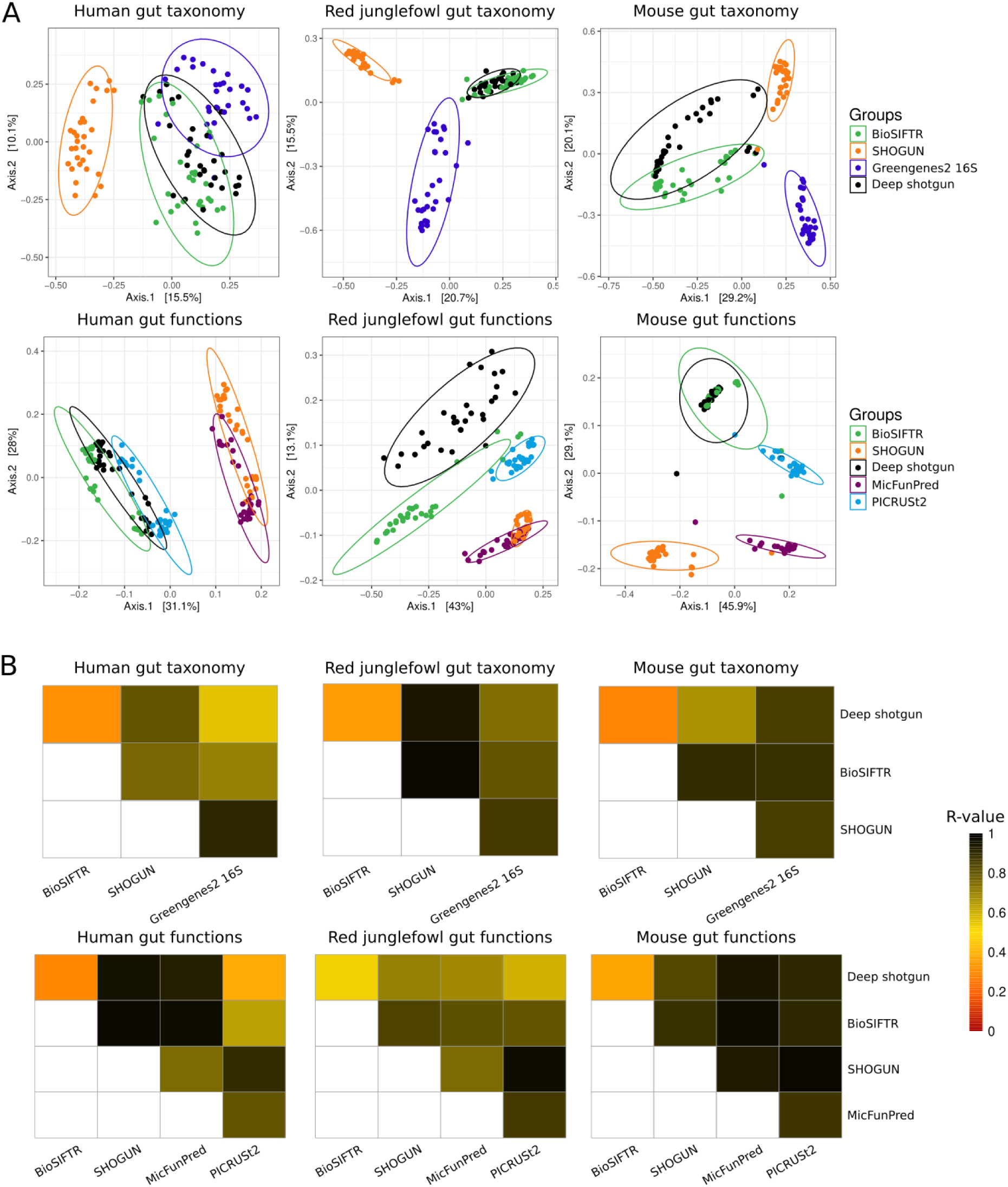
(**A**) Ordination plots of real data profiles using the PCoA method. The dissimilarities between the groups were quantified for taxonomy (top row) and function (bottom row) using Bray-Curtis (for relative abundances) and Jaccard dissimilarity (for presence-absence), respectively. Group size = 30. Ellipse’s confidence = 0.90. (**B**) Heatmaps represent pairwise analysis of similarities (ANOSIM) of feature profiles for real data. The R statistic is depicted in a colour scale; smaller values indicate more similar profiles. All pairwise comparisons were significant (P < 0.01).

These observations support the conclusion that BioSIFTR is the most versatile tool for taxonomic profiling of shallow shotgun data, producing species-level profiles that most closely resemble those obtained from deep shotgun sequencing.

#### KEGG Orthologues annotation

Using deep shotgun sequencing as the gold standard for functional annotation, we observed that data dispersion was homogeneous between deep shotgun and all other tested methods across the three biomes (see Supplementary material, Table S5). Nevertheless, significant compositional differences emerged in the pairwise comparisons with deep shotgun profiles. Consistent with the results from taxonomic profiling, BioSIFTR produced functional profiles most closely aligned with the gold standard across all three biomes (Figures 7 and 8). For the human-gut and red junglefowl-gut datasets, PICRUSt2 ranked second based on the R-value, whereas in the mouse-gut biome, BioSIFTR was the only method that generated a similar profile to the deep shotgun reference. In contrast, MicFunPred and SHOGUN consistently produced the most divergent functional profiles relative to deep shotgun sequencing in all biomes (Figure 4).

We observed highly similar taxonomic profiles between BioSIFTR and deep shotgun sequencing. However, the functional profiles were less well aligned, suggesting that the chicken-gut database may require the inclusion of additional genomes to improve pangenome coverage and functional annotation accuracy.

Notably, BioSIFTR was the only method that achieved an R-value below 0.5 in all biomes when compared to the gold standard, reinforcing its utility as a robust and versatile tool for functional annotation in shallow shotgun metagenomics.

### 3.5. Real use case in human-gut microbiome and phenotype differentiation

PERMANOVA analysis of samples from Healthy Cohort 1, before and after L-carnitine intervention, revealed no significant differences in either taxonomic or functional microbial community profiles across the tested groups: the overall cohort, vegetarians, and omnivores (see Supplementary Material, Table S6).

In contrast, significant differences (P < 0.01) were observed in the functional profiles of KEGG Orthologs (KOs) and Pfam families between high and low TMAO producers, both in the overall cohort and within the vegetarian and omnivore subgroups. Notably, taxonomic profiles at the species level differed significantly between high and low TMAO producers in the overall cohort but not when vegetarians and omnivores were analysed separately.

However, analysis of group dispersion revealed heterogeneous variance in functional profiles (Supplementary Material, Table S6), indicating that the observed differences may be driven by variation in group dispersion, composition, or both.

Module and pathway visualisation displayed similar metabolic profiles across low and high TMAO producers (Figure 5). For example, TMAO reductase was highly prevalent in both groups. In contrast, the enzyme responsible for converting dimethylamine into monomethylamine was only detected among high-TMAO producers (highlighted in red in Figure 5). However, differential abundance analysis of species profiles and KEGG Orthologs (KOs) did not identify any significantly enriched features.

**Figure 5.**
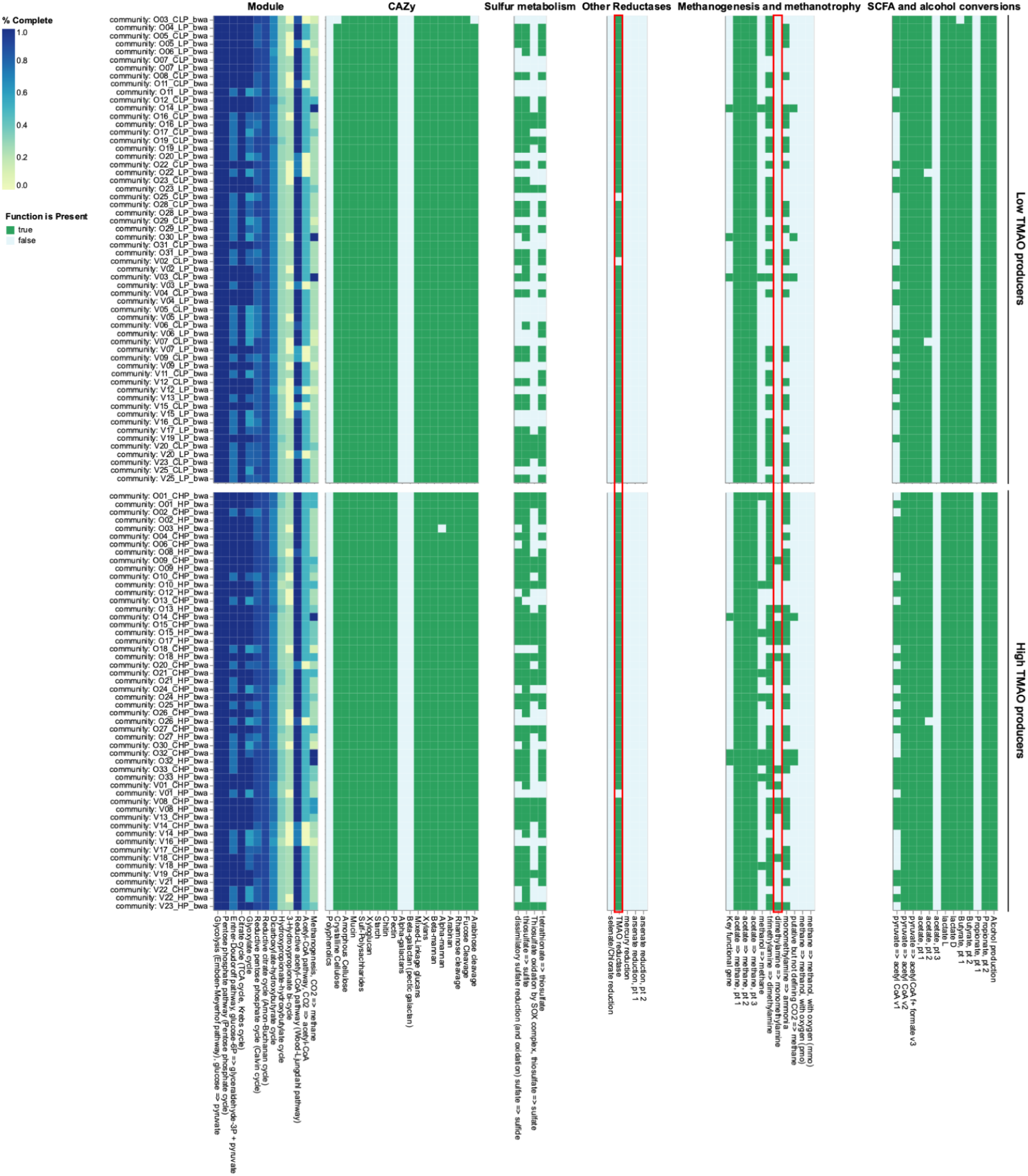
DRAM-style visuals generated with the BioSIFTR pipeline and modified in Vega-Lite v6.1.2 (46) to remove ETC complexes, nitrogen metabolism, and photosynthesis plots. Trimethylamine-related functions are highlighted in red frames.

## Discussion

The BioSIFTR pipeline offers increased resolution and accuracy for functional predictions in microbiome analyses at a sequencing cost comparable to that of 16S rRNA gene amplicon sequencing. By design, through curated biome-specific databases, the method also concentrates on the taxa and functions biologically relevant to each microbial community.

Here, we showed that the functional predictions produced by BioSIFTR using the standard 1 M reads closely match the taxonomic and functional profiles of the complete deep shotgun sequenced metagenomes with more than 20 M reads in the red junglefowl (chicken), human and mouse gut datasets. Furthermore, using the BioSIFTR method at a read depth of 1 M reads per sample, we accurately reproduced the functional potential-related results from a recent study on TMAO production in the human gut microbiome (41). Shallow shotgun sequencing combined with BioSIFTR can thus serve as a high-performance stand-in replacement in studies where functional predictions are currently obtained through 16S rRNA metabarcoding.

16S metabarcoding, combined with PICRUSt2 analysis, is currently one of the most popular low-cost options for functional inference in microbial communities (7). This approach performs well in the human gut biome, but generates divergent functional profiles in other biomes. The other marker-gene-based method we tested, MicFunPred, also generated functional profiles which differed from the deep-sequenced metagenome results. Previous studies have highlighted concerns about the reliability of functional predictions based on 16S rRNA gene amplicon sequencing (7). This is most noticeable when using one or two variable regions, likely due to their limited taxonomic resolution (47). These regions often fail to distinguish between closely related species or strains that may differ significantly in their functional potential (6). As a result, amplicon-based approaches can lead to inaccurate or overly generalised functional predictions.

The main strength of our approach lies in the use of the biome-specific genome databases, which likely reduces the number of false-positive hits to microbes and functions that should be absent from the sample type. Such false positives can significantly impact the conclusions of a microbiome study and potentially lead to pursue further research based on misleading findings (13). While the catalogues are currently prokaryotic-specific, this will change in future versions. Thus, updates to the database could enable the analysis of both prokaryotic and eukaryotic communities together. We advise using BioSIFTR with at least 0.5 M reads per sample in gut biomes, which can be achieved with pre-planned shallow shotgun sequencing. However, our method could also be used when deep shotgun data is low-yield, for instance, due to contamination with host reads. In such cases, researchers have to make the best of the remaining data, e.g., (48). Our method offers an alternative for reanalysing such datasets that otherwise would be challenging to use for functional profiling.

However, the biome-specific catalogues do not currently cover all environments, which limits the applicability of our approach to specific sample types. In particular, more complex biomes such as soil and sediments are underrepresented in the collection, although these catalogues are anticipated to be added to the collection in the near future. Furthermore, the geographical extent of the data is still limited mostly to Western and Westernised countries.

This issue is most prevalent in the human-gut catalogue, where the geographical biases are well-known, e.g., (49). Thus, we advise caution in using our method in samples originating from regions and countries with low-availability of metagenomic sequencing data, such as most low- and middle-income countries. As ongoing sequencing projects expand the geographical scope of microbiome data, these new datasets and genomes can be integrated into MGnify Genomes catalogues through our regular updates. This will help to close the gap in less well-represented geographical regions, e.g., (50). While it might be tempting to use the biome-specific genome databases as a reference for mapping 16S rRNA metabarcoding data, we would advise against that. By virtue of the assembly process used to generate MAGs, the rRNA genes are rarely well represented in MAGs (51). As such these databases are not well-suited for use with 16S metabarcoding data.

Concerns have been raised recently about contamination of MAG databases with human DNA, and the impact it can have on metagenomic studies (13). In the process we describe in the publication, this impact is alleviated in two ways. Where MAGs were generated within MGnify for the catalogues, raw-reads and resulting assembled contigs are mapped against the reference human genome to identify and remove potential human contamination prior to catalogue generation. In addition, the BioSIFTR pipeline includes a quality control step to decontaminate for human, phiX, and host reads. The lower dispersion in the functional profiles generated from deep-shotgun data underlines its robustness as a gold-standard reference. However, deep-shotgun taxonomic profiles are highly dispersed, which hinders comparison with the other methods. While deep-shotgun sequencing remains the most common approach for the functional analysis of metagenomic data, it has limitations associated with data processing and analysis such as the inherent bias associated with assembly of the most abundant taxa, and limited annotation of fragmented genes, and mobile elements, etc. (52, 53). In our opinion, both shallow- and deep-shotgun sequencing have their appropriate applications but as always, users should be aware of the sources of bias and common pitfalls in their interpretation.

## Final remarks

When the BioSIFTR pipeline was applied to real datasets, it generated functional profiles closely resembling the gold standard, even when subsampled to levels as low as 0.1 M reads. This suggests that once a certain threshold of sequencing depth is achieved, profiles become more stable and reliable, as observed by past publications (18, 54). However, the selection of the tool and its optimal settings for taxonomic annotation depends on community richness and is not a trivial process. Gut-associated mammalian samples are generally high in species richness. Nevertheless, there could be cases of low richness as *in vitro* experiments, gnotobiotic specimens, or mock-communities generated in the lab. In general, users cannot be expected to know the true sample richness.

Currently, shallow shotgun sequencing has not yet reached the popularity of 16S rRNA gene amplicon sequencing in the taxonomic characterisation of microbiomes, despite its higher resolution and stability (9). In this study, we demonstrate that the BioSIFTR pipeline infers the functional potential in specific microbiomes from shallow shotgun metagenomic sequencing data. Our approach more closely estimated the functions from deep shotgun metagenomic sequencing than methods that use 16S rRNA gene amplicon sequencing data. Furthermore, we replicated the differences in functional potential of the microbiome in the experimental groups from a previous human gut study, using <2% of the original data. The BioSIFTR method could thus be used as an improved replacement for 16S sequencing in projects where financial constraints prevent the use of deep metagenomic sequencing and functional information would be useful (see e.g., (7)). Such projects could include, e.g., upscaling sampling to increase statistical power (55) or including time-series data to account for temporal effects and variability (56, 57).

The BioSIFTR tool is a powerful approach for functional prediction in specific biomes, which approximates the functional information of a deep shotgun-sequenced metagenome while using only a fraction of the data. We suggest shallow shotgun sequencing combined with BioSIFTR, subject to biome representation in a MAG catalogue, can be a viable alternative for 16S rRNA gene amplicon-based analyses with an increased taxonomic and functional resolution, and lower bias.

## Code and data availability

The BioSIFTR pipeline:

https://github.com/EBI-Metagenomics/BioSIFTR

Methods repo:

https://github.com/EBI-Metagenomics/biosiftr_extended_methods

Synthetic reads for BioSIFTR optimisation are available at the ENA under the project accession PRJEB89162.

## Author Contributions Statement

Conceptualization: A.E-Z., M.O.R., R.D.F. Data curation: J.L. Formal analysis: A.E-Z., M.O.R. Funding acquisition: R.D.F., L.L. Investigation: A.E-Z., M.O.R., L.L. Methodology: A.E-Z., M.O.R., R.D.F., L.R., M.B., L.L. Project administration: R.D.F., L.L. Resources: R.D.F. Software: A.E-Z., M.B. Supervision: R.D.F., L.R., M.B., L.L. Validation: A.E-Z. Visualization: A.E-Z., M.O.R. Writing – original draft: A.E-Z., M.O.R. Writing – review & editing: A.E-Z., M.O.R., M.B., J.L., D.M., L.R., R.D.F., L.L.

## Funding

This project was supported by the European Union’s Horizon 2020 research and innovation programme, the “FindingPheno” project [952914], the Research Council of Finland [338818 to M.O.R.], and the Finnish Cultural Foundation [210944 to M.O.R.]. This work was also supported by the Medical Research Council [grant number MC_PC_21045].

## Acknowledgements

The authors wish to acknowledge CSC – IT Center for Science, Finland, for computational resources.

## Conflict of Interest

None declared.

## Notes

https://github.com/EBI-Metagenomics/BioSIFTR/

https://github.com/EBI-Metagenomics/biosiftr_extended_methods

### Competing Interest Statement

The authors have declared no competing interest.

