## Supplementary figures and images for "Biome-specific genome catalogues reveal functional potential of shallow sequencing"

### S1_biosiftr_optimisation.pdf

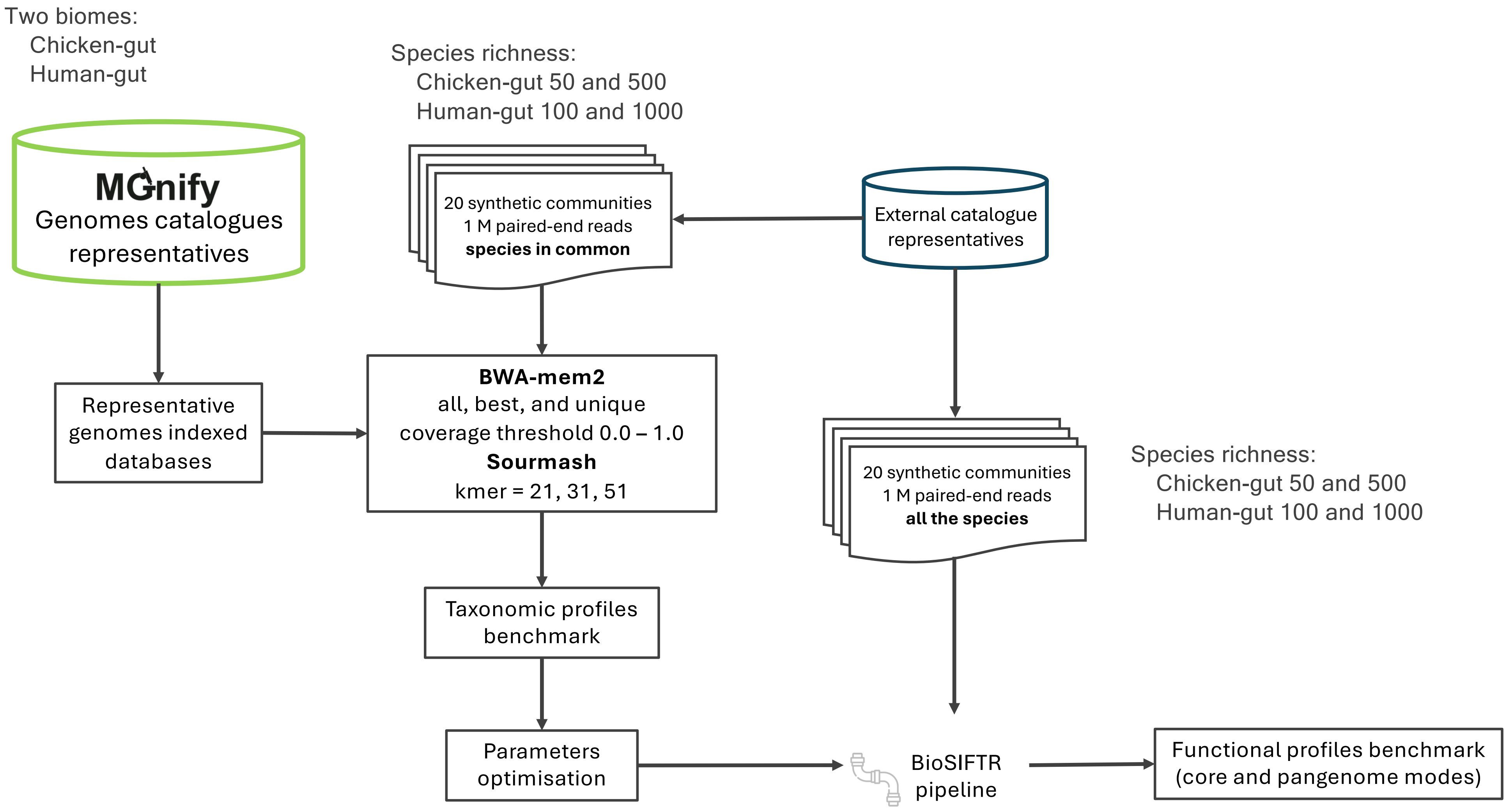

### S2_real_data_comparison.pdf

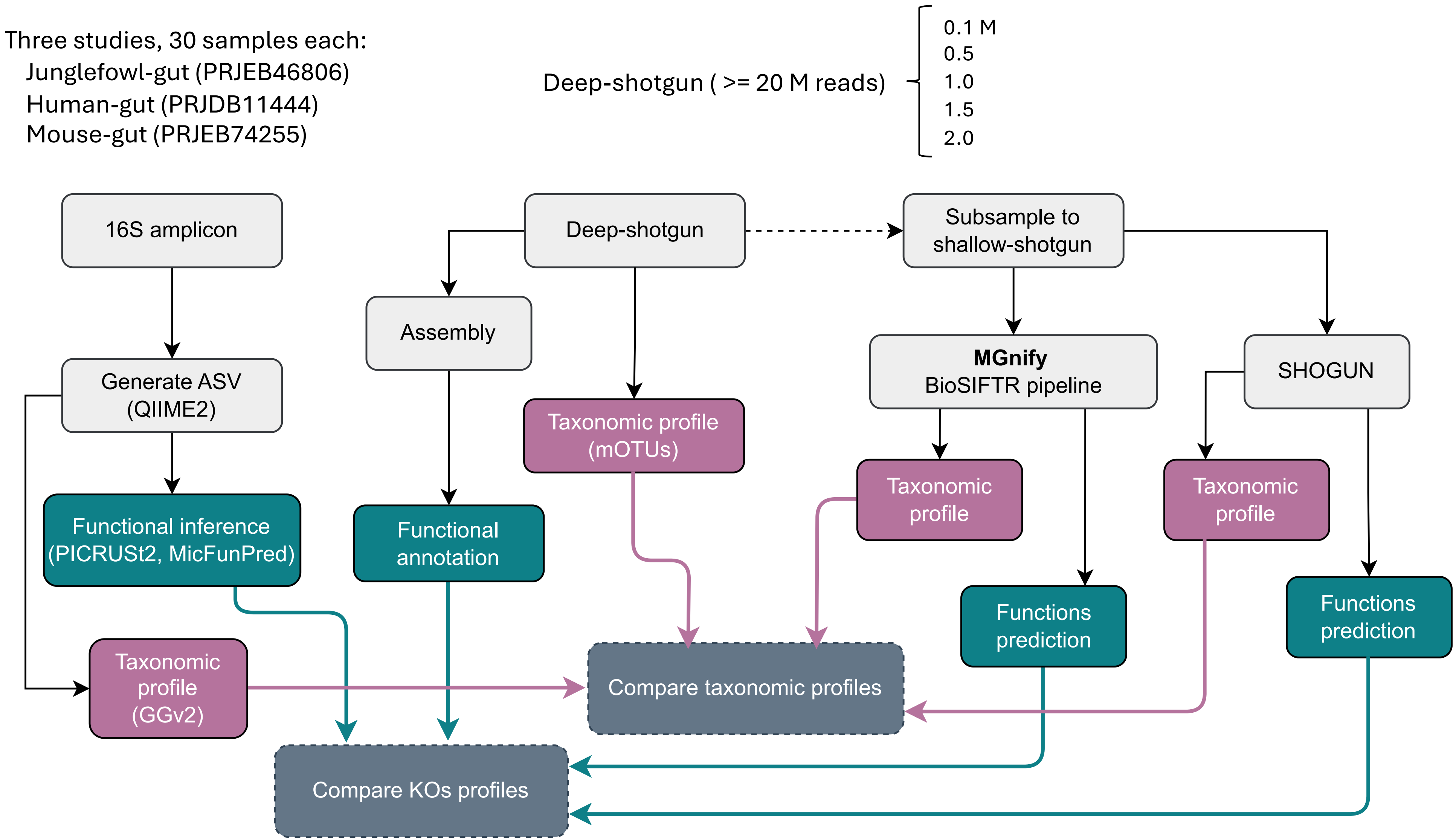

### visual_abstract_full.pdf

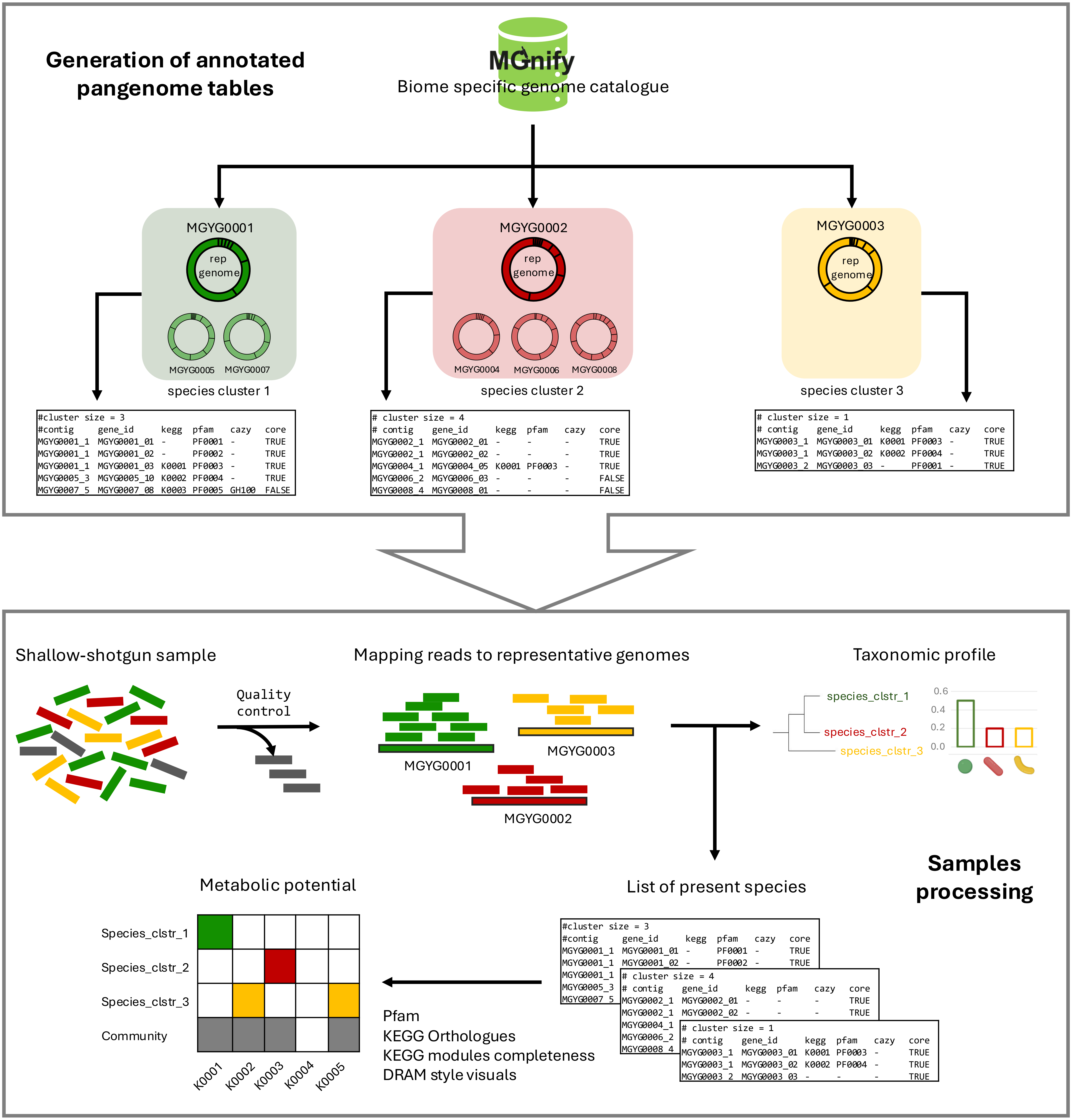
